# Comparative Dual RNA-Seq Analysis of Eight *Orientia tsutsugamushi* Strains in Endothelial Cells Reveals Regulatory Patterns in Bacterial and Host Pathways During Infection

**DOI:** 10.1101/2025.12.01.691569

**Authors:** Regan Hayward, Suthida Chuenklin, Chitrasak Kullapanich, Panjaporn Chaichana, Brian Ferguson, Lars Barquist, Jeanne Salje

## Abstract

*Orientia tsutsugamushi* is an obligate intracellular bacterium that causes the mite-borne human disease scrub typhus. The species is characterised by having many strains that differ in their ability to cause disease in murine infection models and in human patients. The genomes of a diverse set of *Orientia tsutsugamushi* strains have recently been analysed in detail, revealing broadly similar gene content with genomes differing primarily in synteny and in the composition of a large arsenal of predicted secreted effector proteins. Given the similarity in gene content, here we asked whether the observed differences in virulence were driven by genome-wide differences in gene expression of bacterial genes, and differences in the ensuing response of infected host cells. To explore this, we carried out a dual RNA sequencing analysis of eight diverse strains of *Orientia tsutsugamushi* grown in cultured human endothelial cells. An overall analysis found no clear patterns in the bacterial or host gene expression patterns that correlate with the ability to cause disease. We found that all strains induce a strong type 1 interferon response in endothelial cells, but that within that broad response each strain has a unique fingerprint of induced genes likely leading to different disease outcomes when combined with a complex immune system *in vivo*. We compared expression levels of orthologous genes between different strains and identified some bacterial pathways with constant expression levels whilst others were more variable, leading to the identification of specific pathways under tight transcriptional control during cellular infection. Together our data show that inter-strain differences in virulence are not defined by expression levels of any one set of bacterial or host genes and also reveal insights into transcriptional regulation of different pathways in *Orientia tsutsugamushi*.

## Introduction

*Orientia tsutsugamushi* (Ot) is an obligate intracellular, Gram negative, Alphaproteobacterium in the Order Rickettsiales, that causes the mite-borne human disease scrub typhus^1^. Symptoms include headache, fever, rash and myalgia, and if effective antibiotics are not promptly administered it can escalate to multiple organ failure and death, with a median mortality of 6% in untreated cases^2^. Scrub typhus caused by Ot is a leading cause of severe febrile illness in many parts of Asia, especially in rural settings^3,4^. The known global distribution of scrub typhus was recently expanded with the discovery of two new species: *Candidatus* Orientia chiloeensis and *Candidatus* Orientia chuto, in Latin America and the Middle East respectively^5–8^.

Ot is an obligate intracellular bacterium that replicates directly in the cytoplasm of infected cells^9^. As a mite-transmitted bacterium, it initially grows in the skin at the site of inoculation, forming a painless necrotic lesion called an eschar. The bacterium disseminates from here through the blood and lymphatic systems to lymph nodes and multiple organs including lungs, liver, spleen, brain, kidneys, and heart^10–12^. The primary *in vivo* target cells of Ot are dendritic cells, macrophages, and endothelial cells, although growth in other cell types has been demonstrated *in vitro* including in fibroblasts and epithelial cells^13–17^. The intracellular lifecycle of Ot has been well studied *in vitro.* Bacteria enter cells through clathrin-mediated endocytosis and macropinocytosis^18,19^, then escape from late endosomes^18^. They traffic along microtubules to the perinuclear region where they undergo replication in a tightly packed bacterial microcolony^20^. Bacteria then move to the edge of infected cells and exit through a budding mechanism, in which they are encased in host plasma membrane as they leave the infected host cell^11^. We recently showed that the bacteria that have budded off the surface of infected cells are in a different developmental stage, called the extracellular bacteria (EB) stage, compared with their replicating intracellular bacteria (IB) counterparts^19^.

We previously sought to examine the interplay between Ot and host cells by carrying out a dual RNA sequencing (dual RNAseq) study using two Ot strains, Karp and UT176, grown in human umbilical vein endothelial cells (HUVECs)^21^. We showed that both strains induced a type I interferon antiviral response, but there were differences in the cytokines induced by the two strains. Since UT176 was found to be less virulent in a mouse model than Karp we concluded that these differences may lead to differences in disease outcome. One limitation of that study is that we only compared two Ot strains.

Ot has an unusual genome, which is dominated by a highly amplified integrative and conjugative element called the Rickettsiales amplified genetic element (RAGE)^22,23^. This RAGE has proliferated to such a degree that it now comprises around 50% of the Ot genome. In addition to the RAGE, Ot encodes numerous DNA transposons, retrotransposons, and a gene transfer agent^24^. Together these mobile genetic elements have had an impact on the evolution of different strains of Ot. A recent comparison of the complete genomes of eight Ot genomes showed that whilst these strains were closely related in terms of genes present in the inter-RAGE regions, and shared a 16s rRNA sequence identity of >99%, the genomes completely lacked synteny with respect to one another, with conserved inter-RAGE regions being shuffled into different positions in different strains, together with variability in the number and composition of the RAGE elements^24,25^. Importantly, the strains differed in the number and identity of RAGE-encoded secreted effector proteins, namely the Ankyrin repeat containing family of proteins (Anks). Given the unusual variation in gene order and RAGE composition between otherwise closely related strains of Ot, we set about to extend our previous study by carrying out a dual RNAseq study of eight strains of the species Ot.

In this study we carried out dual RNAseq analysis of seven strains of Ot (Karp, Kato, Gilliam, UT76, Ikeda, TA686, TA763) and compared it with our previous analysis of Karp and UT176^21^. These were grown in HUVEC cells as an established model of endothelial cells. By carrying out a large comparison using a total of eight different strains we could identify robustly conserved host pathways upregulated or downregulated in response to all strains of Ot, as well as pathways that were only affected in response to certain strains. We also compared the expression levels of bacterial genes in eight strains of Ot and identified bacterial gene expression networks with constant or variable expression between strains. Together these analyses allow us to make predictions about the role and regulation of different pathways during infection.

## Results

### Host and bacterial transcriptomes do not cluster according to virulence or phylogeny

We performed dual RNA-seq on seven different Orientia strains: Gilliam, Ikeda, Karp, Kato, TA686, TA763 and UT76, that were originally isolated from humans and small rodents (Supp. Table 1). Human umbilical vein endothelial cells (HUVEC) were infected and grown for 5 days using 3 separate biological replicates (Supp. Table 2) as per our previous study^21^. Post-infection, we carried out rRNA depletion and sequencing of the samples, generating between 55-75 million reads (Supp. Table 3). To accurately quantify host and bacterial reads, particularly those originating from the repeat-rich regions typical of *Orientia* genomes, we used the dualrnaseq pipeline with Salmon selective alignment.

We observed a wide range in the proportion of bacterial reads in infected HUVECs across strains, ranging from 4% with TA763 to 57% with TA686 (Fig. 1A, Supp. Table 4). Examining the distribution of reads to RNA classes shows that rRNA depletion was successful for both the host (<0.1% rRNA reads) and each bacterial strain (<3% rRNA reads), resulting in the majority of reads being assigned to coding sequences (CDS), facilitating downstream analyses (Fig. 1B-C, Supp. Table 4).

**Figure 1:**
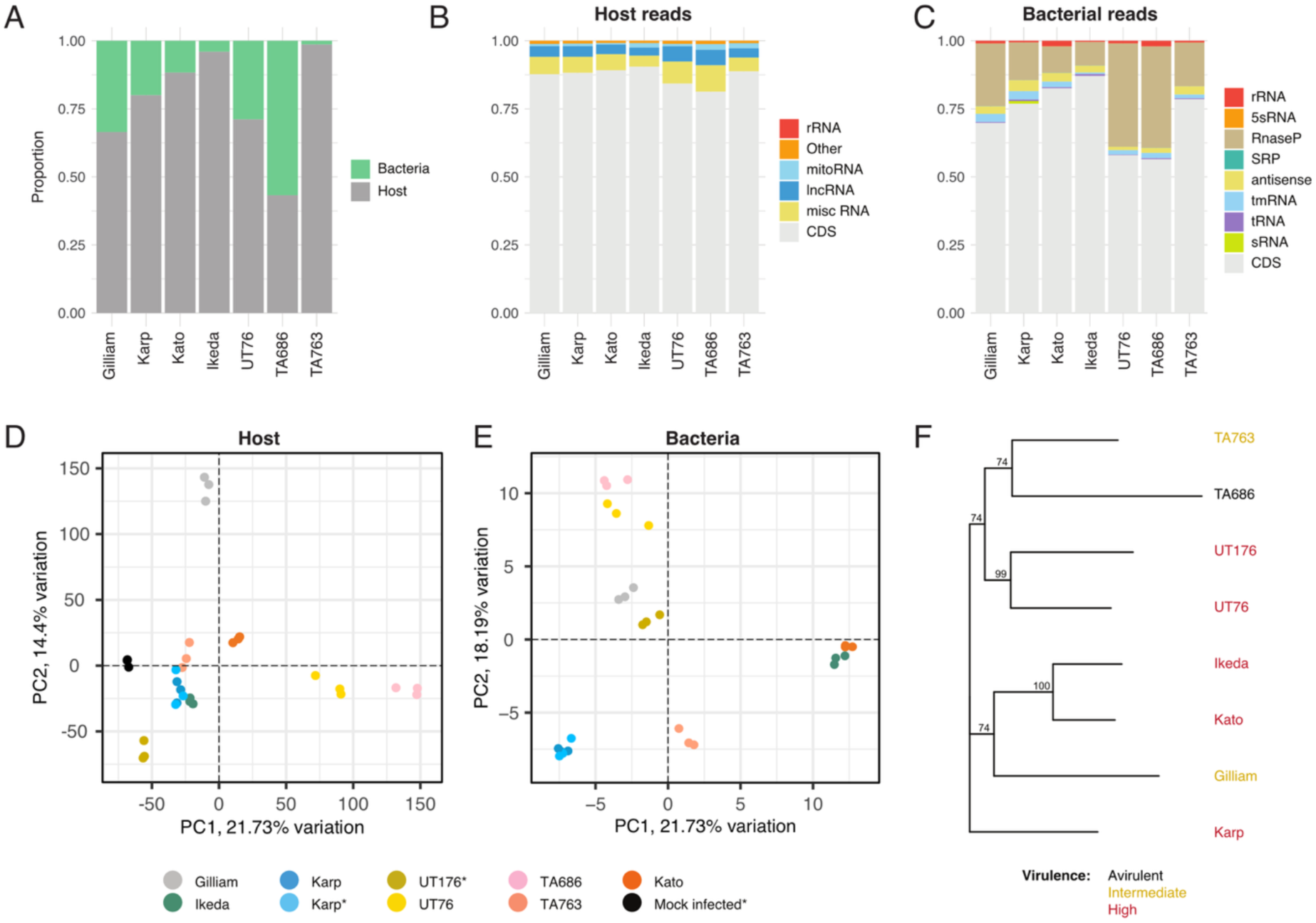
Dual RNA-seq of infected HUVEC cells from different Orientia strains. A. Proportions of host and bacterial reads for each Orientia strain, averaged over three replicates. B. Proportions of reads assigned to different RNA classes in HUVECs (human umbilical vein endothelial cells) and C. across each Orientia strain. D. PCA (Principal component analysis) bi-plot of host reads. E. PCA bi-plot for orthologous bacterial reads. The star (*) indicates additional data used from our prior study. F. Phylogenetic tree of the eight Orientia strains, colour-coded by virulence levels^26^.

In assessing the consistency across biological replicates for each strain, we separately analysed the correlations of host and bacterial reads. We observed high correlations in both host (0.85-0.88) and bacterial (0.92-0.99) reads across strains, further supporting the robustness of the underlying data (Supp. Table 4).

The experimental procedures from this study mirrored the procedures from Mika-Gospodorz etc al.^21^, providing an opportunity to incorporate additional RNA-seq data for a more comprehensive analysis. This allowed us to include an additional Ot strain UT176, three additional Karp replicates and mock-infected HUVEC cells for host comparisons. We used RUV-seq to remove batch effects between datasets, which resulted in high similarity in the now six Karp replicates with principal component analysis (PCA) (Fig. 1D).

Using the 8 strains, we identified 706 orthologous bacterial genes present in all genomes in our study, which we used for PCA. The resulting bi-plot also showed high similarity in Karp replicates, but different clustering compared to the host PCA (Fig. 1E). The relative virulence of these strains has been determined based on a combination of murine infection studies and prevalence in human clinical cases^26^. UT76, Karp, UT176, Kato and Ikeda are high virulence strains, with Kato and Ikeda leading to the most severe outcomes in a murine infection model. TA763 and Gilliam are intermediate in virulence, causing disease in humans but with limited virulence in murine models that, in that case of Gilliam, is dependent on the mouse strain. TA686 is avirulent and has not been isolated from human patients and does not cause disease in a murine infection model. Surprisingly, both host and bacterial PCA did not show any grouping related to virulence or phylogeny (Fig. 1F)^26^.

### All Ot strains elicit a strong Type I interferon response, but induction of downstream genes differs between strains

We explored the inflammatory response to infection of endothelial cells by Ot (Fig. 2A). All strains induced a strong type 1 interferon response, as has been reported extensively previously^21,27–31,32^ with pathway analysis highlighting a viral-like inflammatory response as well as the induction of specific pathways linked to RIG-I-like signalling, cytosolic DNA sensing, NOD-like signalling, and Toll-like signalling. There was significant overlap between these pathways and the different pattern recognition receptors (PRRs) previously implicated in Ot recognition: the toll-like receptor TLR2^33^, the lectin Mincle^34^, the peptidoglycan receptors NOD 1/2^17,35^, and cytosolic DNA and RNA sensors cGAS/STING and RIG-I^32^ (Fig. 2B). We further explored the relative roles of these different PRRs in the response to different Ot strains by determining the differential expression of each of these PRRs for each strain (Fig. 2C, Supp. Fig. 1). PRR induction was generally low, as expected for receptor genes. Gilliam stands out as generally inducing a weaker response than the other strains in our analysis. This is particularly notable in the case of *NOD2*, in which all the strains cause an increase in transcription with the exception of Gilliam that results in a decrease in receptor expression.

**Figure 2:**
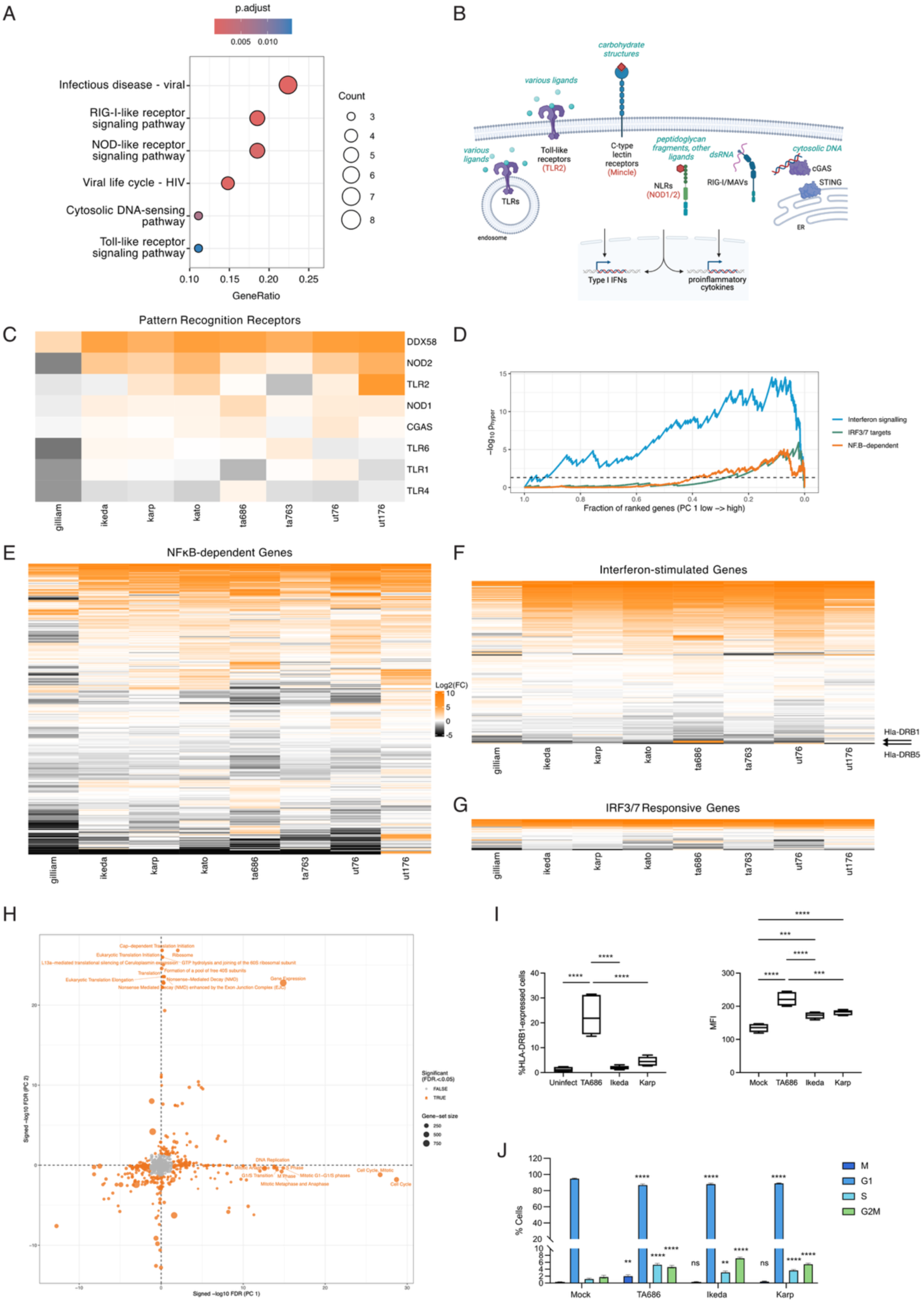
An overview of the host response to different Ot strains. A. Pathway analysis showing the pathways induced by multiple Ot strains. The size of the circle indicates the number of strains (1-8) that induce this pathway, and the color indicates the adjusted p-value. B. Schematic illustration showing the different pattern recognition receptors previously implicated in detection of intracellular Ot. C. Heatmap showing the log_2_ fold-changes (logFC) between infected and mock-infected cell for genes involved in TLR, C-type lectin, RIG-I-like and NOD-like receptors. Larger version of this heatmap containing gene names is given in Supp. Fig. 1. D. Cumulative hypergeometric enrichment of immune-response genes enriched along PC1 in the PCA analysis shown in Fig. 1D. E-G. Heatmaps showing the log_2_ fold-changes (logFC) between infected and mock-infected cell for genes responsive to NFkB, interferon and IRF3/7. Larger versions of these heatmaps containing gene names is given in Supp. Figs. 2-4. H. A Wilcoxon rank-sum test was used to test the signed loadings within each gene set to those outside it for a difference in median value. Axes show the –log_10_ P-value from the test, while bubble sizes show the gene set size. A version of this figure with more points labelled is given in Supp. Fig. 5. I. Quantification of HLA-DRB1 expression in HUVEC cells infected with TA686, Ikeda or Karp, showing percentage of cells with HLA-DRB1 signal and mean fluorescence intensity (MFI). Statistical significance determined using a one-way ANOVA with Tukey’s multiple comparisons. HLA-DRB1 levels are higher in cells infected with TA686 than uninfected cells or cells infected with Karp or Ikeda. J. Cell cycle distribution of HUVEC cells infected with TA686, Ikeda or Karp compared with uninfected cells. Graph shows mean and standard deviation. Statistical significance determined using a two-way ANOVA with Dunnett’s multiple comparisons, with comparisons to uninfected shown. ** p ≤ 0.01, *** p ≤ 0.001, *** p ≤ 0.0001.

Next, we broadened out our analysis to explore a full set of interferon-, IRF 3/7- and NF-κB-responsive genes to determine the responses to type 1 interferons, signalling through TLR/NOD/lectins, and signalling through cGAS/STING respectively. We asked whether differential induction of these pathways accounted for the variation in host response between strains. We analysed the PC1 component of the PCA analysis shown in Fig. 1D and found that interferon signalling, IRF3/7 targets and NF-κB-dependent genes contributed substantially to the variance along this axis (Fig. 2D). Heat maps showing differential expression of specific genes in these pathways revealed that overall patterns of response were similar between strains, but there was substantial variation in the induction of individual genes (Fig. 2E-G and Supp. Fig. 2-4). These did not follow a simple pattern in terms of virulence or phylogeny and demonstrated that each strain induces a unique fingerprint of inflammatory response that drives differential disease outcome. This is consistent with the conclusions of a recent report from our group^26^.

The avirulent strain TA686 was unique in causing an increase in expression of the interferon-stimulated genes *hla-drb1* and *hla-drb5,* components of the major histocompatibility complex class II (MHC-II) (Fig. 2F and Supp. Fig. 2). These genes were downregulated in all other strains in our study, reminiscent of recent reports that Ot strain Ikeda leads to a decrease in MHC-I expression in infected cells^36,37^. We sought to confirm this finding by measuring the relative levels of DRB-1 protein in endothelial cells infected with TA686 or Karp, compared with uninfected cells, by flow cytometry analysis (Fig. 2I). We found that TA686 caused increased expression of HLA-DRB1 compared with uninfected cells or those infected with Karp or Ikeda, consistent with the observed transcriptional upregulation.

Further analysis of the PCA analysis revealed groups of genes that drove variation in the host response to different strains in PC1 and PC2 (Fig. 2H). Variation in PC1, which was dominated by TA686 and, to a lesser extent, UT176 (Fig. 1D, Supp. Fig. 6) included gene groups in cell cycle and mitotic phase processes. By contrast variation in PC2, driven by Gilliam (Fig. 1D) involved groups of genes carrying out translation, ribosome and gene expression. To explore this pattern further we explored the relationship between infection and host cell cycle in cells infected with Karp, Ikeda and TA686, compared with uninfected cells (Fig. 2J). We found that all three strains caused a change in cell cycle distribution, consistent with previous reports of S-phase block in Ikeda-infected cells^38^. TA686 caused an increase in M phase cells that was not seen in cells infected with Karp or Ikeda. This is consistent with an upregulation of G2/M genes in TA686-infected cells (Supp. Fig. 6) and may reflect a difference in the impact on the cell cycle between different Ot strains.

### A comparison of transcript levels across eight Ot strains reveals pathways with tightly regulated expression during cellular infection

There is evidence suggesting that Ot undergoes a limited developmental cycle during cellular infection^19^, but the genes required at different bacterial stages of growth are not well described. Despite using the same MOI and infection time for all strains, small differences in growth rate and conditions means that cultures will be asynchronous by the time the samples were harvested at 5 days post infection. We exploited this fact to compare the relative expression levels of orthologous genes across the diverse Ot strains, to identify pathways with tightly regulated expression levels at this stage of the intracellular infection cycle. Genes with similar expression levels in all strains are likely constitutively expressed during these growth conditions, whilst large fluctuations between strains reflects pathways whose expression may reflect strain-specific differences in biology, or alternatively may be regulated as part of the developmental cycle.

To identify variably expressed genes across strains, we developed a regression-based method to find genes with higher-than-expected variance given their expression level (Figure 3A; Methods). With a conservative threshold, this identified 70 genes with highly variable expression across strains. Analysis of the heatmap of 70 highly variable genes revealed that there were no clear groupings of genes or strains (Fig. 3B). A detailed analysis of the expression in each strain of the ten most variable genes is shown in Fig. 3C. A range of different genes were present in the list of 70 variable genes, including 31 genes of unknown function, three Ankyrin repeat containing proteins, and the surface protein ScaA. Pathway analyses revealed that these genes did not group into any particular functional category. The observed highly variable expression of these 70 genes suggests that they are under differential regulatory control during intracellular growth in endothelial cells. Similarly, the genes with the lowest variation in expression level between strains were identified (Fig. 3D). When selecting for those with high expression, this notably included genes encoding core components of the RNA polymerase (*rpoB*, *rpoC*), a subunit of DNA gyrase (*gyrB*), translation initiation and elongation factors (*infB*, *fusA*), and an iron-sulfur cluster assembly protein (*iscC*). These genes are likely not subject to significant differential transcriptional regulation under these growth conditions, and may serve as valuable housekeeping genes in future studies.

**Figure 3:**
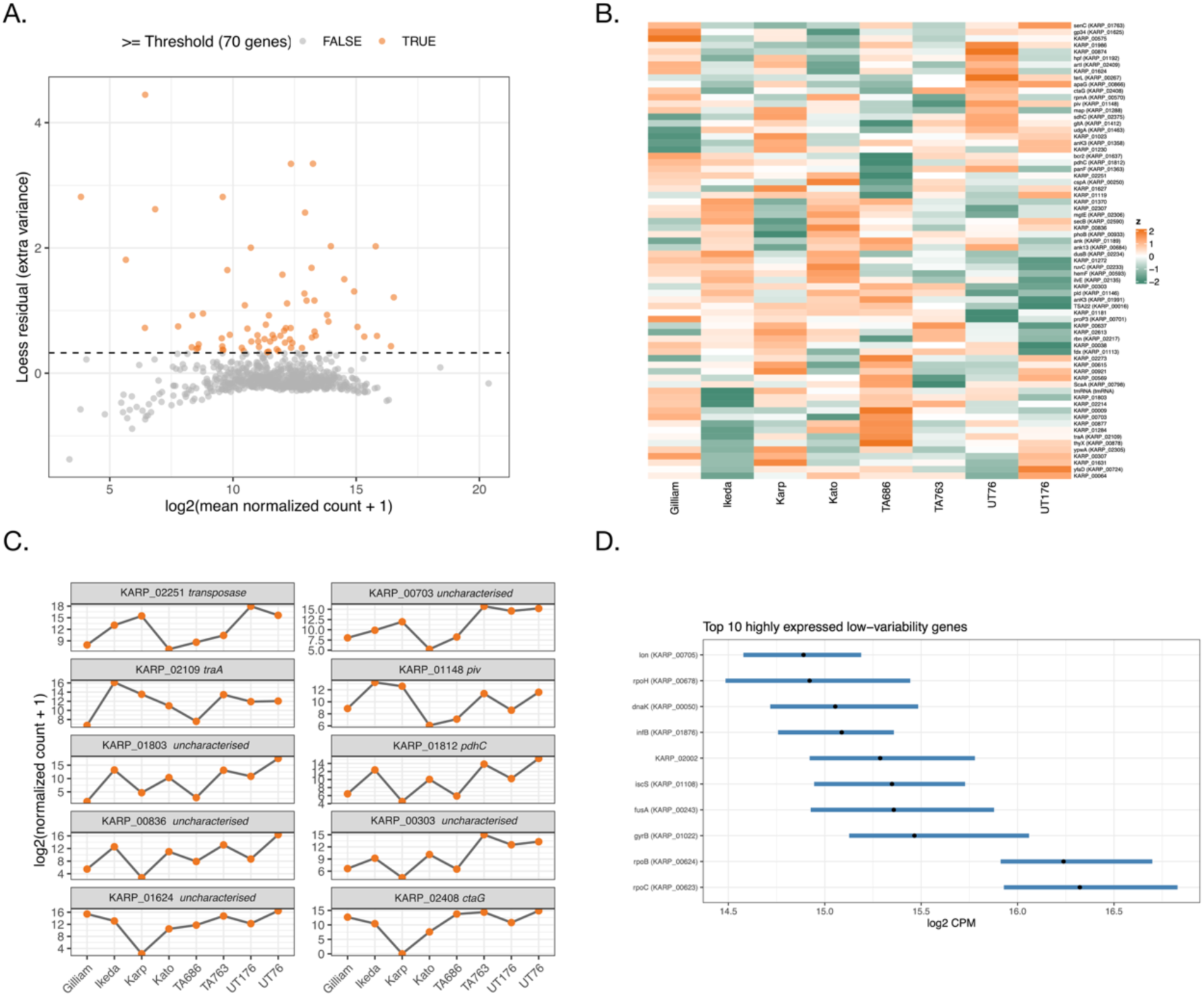
Expression variability across different Orientia strains. A. Scatter plot showing the relationship between gene expression (log_2_ mean normalized counts) and residual variance compared to a fitted LOESS curve. Genes highlighted in orange exhibit a residual variance greater than three times the median absolute deviation of the residual variances across all genes. B. Row-clustered heatmap illustrating the relative expression of 70 genes identified as exhibiting excess variability across all strains. C. Absolute expression profiles of the top ten most variable genes across strains. D. Variation in gene expression in the least variable genes across all strains.

### Expression of bacterial surface proteins, secretion machinery, and secreted effectors, differs between Ot strains

The differential response of host cells to bacterial infection, described above, is expected to be driven in large part by the different bacterial proteins that the host cell encounters. As a cytoplasmic bacterium lacking an LPS layer, the proteins embedded in the bacterial outer membrane play a key role in interactions with host cell machineries. Similarly, bacterial effector proteins secreted from the bacterium into the host cell cytoplasm play an important role in the ability of bacteria to manipulate the host cell response to infection. In this section, we analysed the relative expression of bacterial proteins that are involved in direct interactions with host cells.

We compared the relative expression of Ankyrin repeat containing proteins (Anks). The Ank repeat is one of the most common protein motifs found in nature and mediates diverse protein-protein interactions in eukaryotic cells^39^. However, it is rarely found in bacteria, being primarily found in pathogens that have exploited the fold in their secreted effector proteins that interact with and block host cell protein activities^40^. Ot encodes an enormous and diverse arsenal of Ank effector proteins, with seven conserved Anks (Ank03, Ank08, Ank09, Ank10, Ank11, Ank12, Ank20) present in all described strains, sometimes in multiple paralogous copies^24^. There are an additional 122 diverse Ank orthologs encoded across Ot strains, with a subset of 47-66 present in any one strain^24^. A comparison of the differential expression of the conserved orthologous groups as well as the less conserved Anks revealed substantial diversity between strains (Fig 4A, B). Whilst the exact functions of each of these Anks has not been described, they have been shown to traffic to different cellular compartments in strain Ikeda^41^, and several Ikeda Anks have been shown to perturb specific host cell pathways^38,42–45^. Therefore it is likely that the differential presence and expression of Anks is an important driver of the observed differences in host transcriptional response and virulence^46^.

Next, we compared the expression of surface proteins between Ot strains. Given the small genome of Ot there are a limited number of predicted outer membrane proteins in the Ot genome and these are shown in Fig 4C. Note that *tsa47* (type surface antigen 47)^47^ is predicted to encode a periplasmic serine protease and is thus not an outer membrane protein however it is a highly immunogenic bacterial protein and is therefore included in this analysis. TSA22, TSA47 and TSA56 are bacterial proteins that exhibit strong immunogenicity in human patients^48^. TSA56 is the most abundant bacterial surface protein in Ot and has been shown to play a role in bacterial attachment and invasion into host cells^49^. Whilst *tsa47* was present at similar levels across strains, levels of *tsa22* were comparatively lower in UT176 in a similar pattern to *tsa56.* In addition to the TSA proteins, that are similarly named based on their immunogenicity but share no structural homology, Ot encodes a group of outer membrane autotransporter proteins that all share a conserved beta barrel membrane domain^50,51^. They vary in their soluble passenger domains that extrude into the host cell cytoplasm. Surface cell autotransporter (Sca) proteins are common in host associated bacteria and commonly mediate interactions with host cells^52^. ScaA and ScaC have been shown to be involved in bacterial adhesion^51,53^, whilst ScaC has recently been shown to also mediate bacterial trafficking along microtubules by activating the dynein adapters BicD1 and BicD2^54^. The functions of ScaD and ScaE are unknown. Here we found that *scaA* was the most highly expressed *sca* gene, in addition to being a highly variable gene identified in the previous section (Fig. 3B) with *scaC, scaD* and *scaE* exhibiting comparatively low expression. In addition to transcriptional regulation, it is known that autotransporter proteins often regulate activity through cleavage of the passenger domain^52^, a level of posttranslational regulation that would not be detected in the current analysis.

In addition to Anks, Ot encodes a diverse arsenal of tetratricopeptide repeat containing proteins (TPRs). This is also a common eukaryotic fold mediating protein-protein interactions^55^ and also exploited by pathogens^56^. The TPRs of Ot are less well characterized than the Anks, but are thought to be secreted into the host cytoplasm and mediate protein-protein interactions^57^. TPRs are more difficult to classify than the Ot Anks^24^ and therefore we compared the bulk expression levels of 21- 48 TPRS encoded by each strain (Fig. 4D). The strains exhibited different levels and distributions of TPR proteins, with UT176 exhibiting the lowest overall TPR expression and Kato and TA686 the highest. Since Kato is the most virulent strain in our collection and TA686 is avirulent^26^, these patterns do not correlate with virulence.

Ot encodes two Type 4 secretion systems (T4SS): a *vir*-type T4SS present in a single copy dispersed across six locations in the genome, and an F-type T4SS present in the multi-copy integrative and conjugative RAGE of Ot^24^. The F-T4SS is abundant, with 370-604 copies of its genes in the different Ot strains^24^. The activity and cargo of both systems are not well established. We compared the expression of T4SS genes across the Ot strains (Fig. 4E). Given the similarity between the *vir* and F-T4SS systems, there are a number of genes with identical sequences present in both systems, making read assignment difficult. We therefore analyzed expression of the *vir* and F-T4SS systems both including (Fig. 4E) and excluding (Supp. Fig. 7) duplicated genes. Both analyses showed that *vir* transcripts were more abundant on average than T4SS transcripts, and that the levels of expression were generally similar across different Ot strains.

### Analysis of bacterial growth genes reveals that certain genes involved in peptidoglycan polymerisation and recycling are transcriptionally regulated

The regulation of molecular machines that carry out bacterial growth and division in Ot is poorly understood. We sought to use the current dataset to explore different aspects of bacterial growth and division and identify the components of these pathways that are differentially expressed between strains, suggesting either a role in bacterial differentiation or a different function between strains. Peptidoglycan is the molecular mesh that encases bacterial cells and comprises the cell wall^58,59^. Bacteria encode two large multiprotein complexes, the divisome and elongasome (Fig. 5A)^60–62^, that ultimately function to localize peptidoglycan polymerization machines spatially and temporarily, enabling tight regulation of growth and division through regulation of the cell wall.

**Figure 4:**
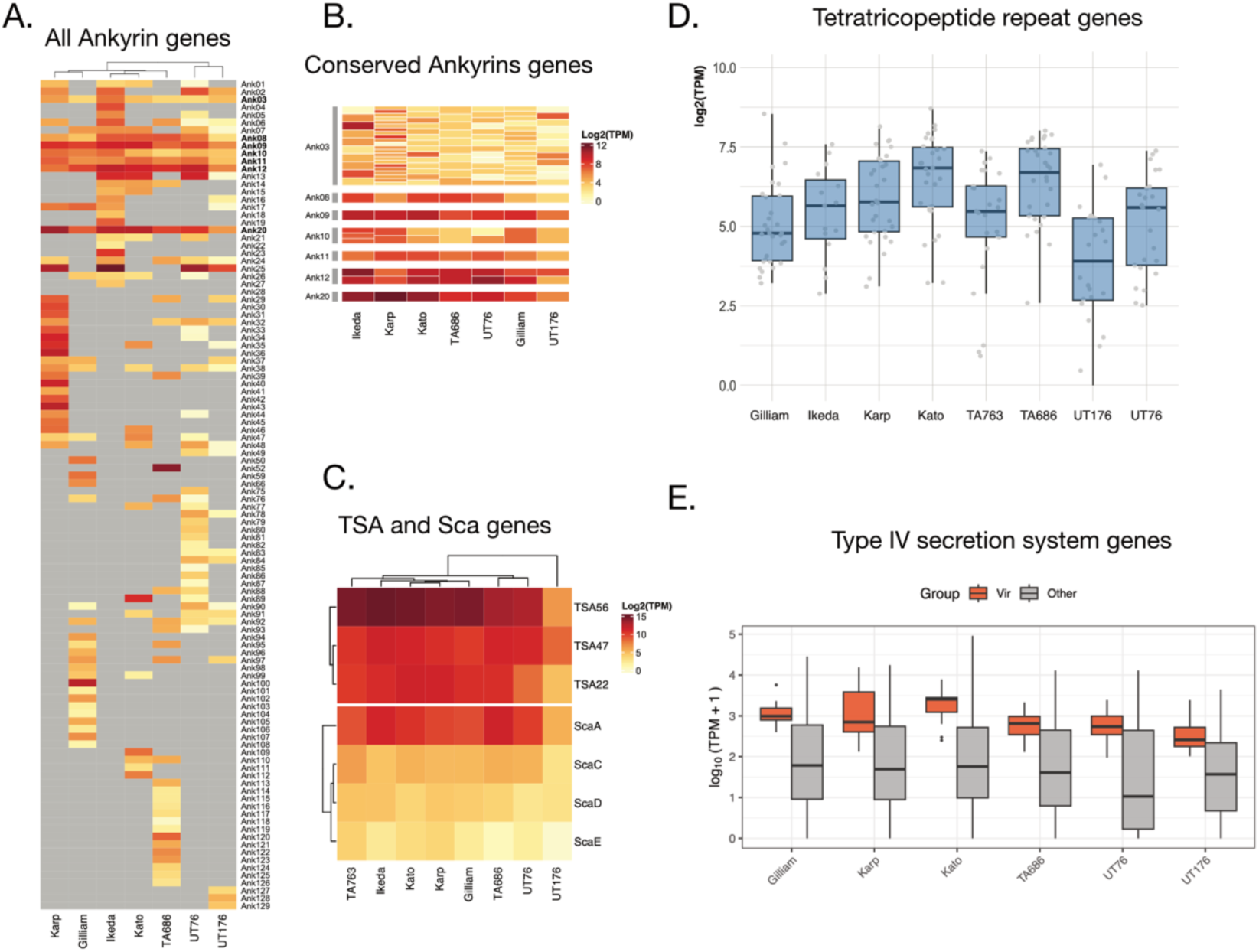
Ot infection requires manipulation of the host response. A, B. Heat maps showing the relative expression of all (A) and conserved (B) Ankyrin repeat containing genes in seven strains of Ot. Ankyrins conserved across all strains in this study are shown in bold in (A). C. Heat map showing the relative expression of genes encoding surface proteins and major antigens in eight strains of Ot. D. Boxplot showing the expression levels of all 21- 48 TPR genes in each of eight strains of Ot. E. Boxplot showing the expression levels of *vir-*T4SS and non-*vir-*T4SS genes in Ot. The non-*vir* genes primarily belong to the F-type T4SS of Ot but cannot all be definitively classified as such due to duplicated genes shared between *vir-* and non-*vir-*T4SSs. A version of this analysis excluding all duplicated genes is given in Supp. Fig. 7.

**Figure 5:**
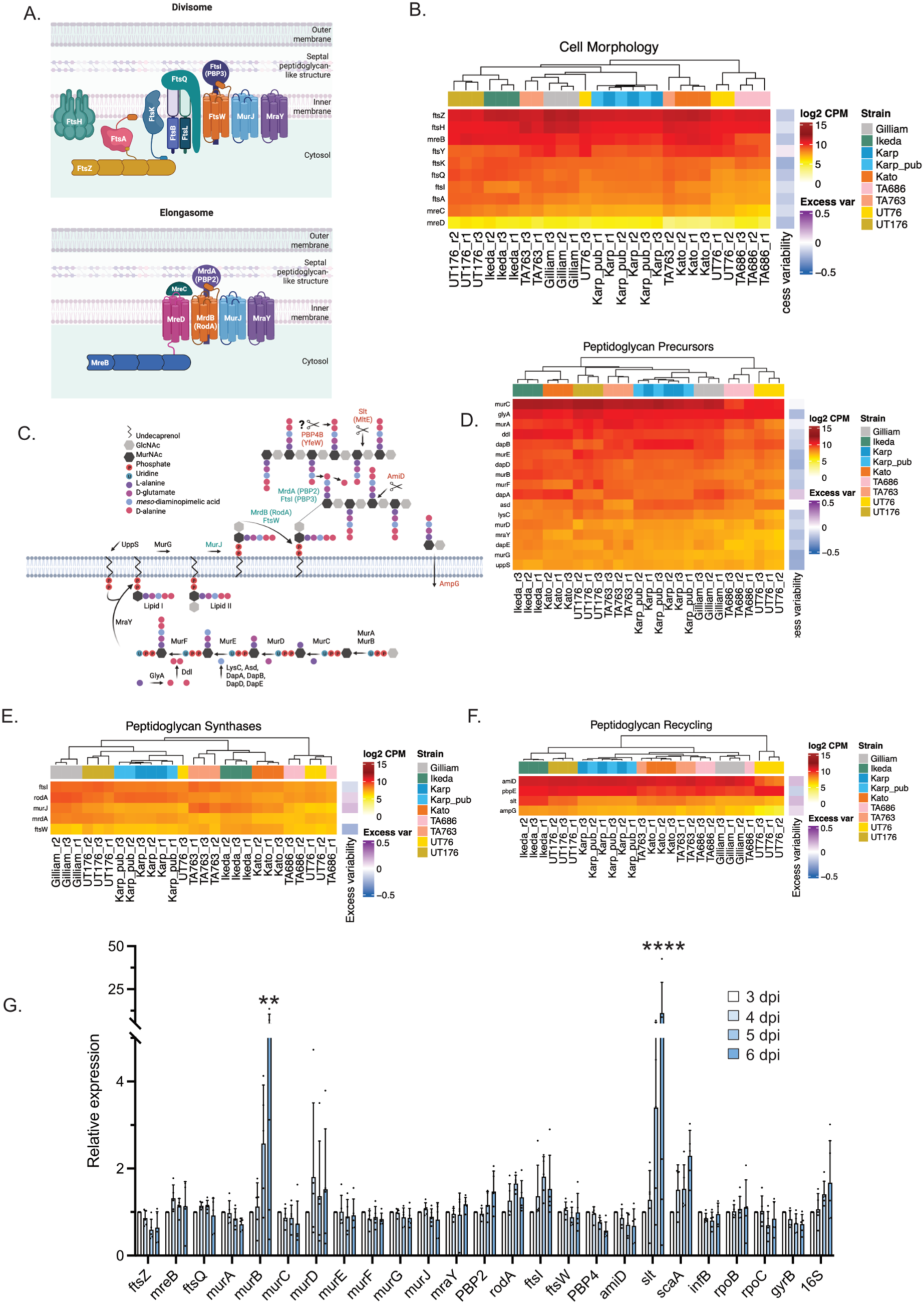
Expression of peptidoglycan-like genes across 8 Orientia strains. A. Schematic diagram showing components of the divisome and elongasome machinery in Ot. B. Heat map showing expression of cell morphogenesis genes in eight strains of Ot. C. Schematic diagram showing the peptidoglycan biosynthesis pathway in Ot. D-F. Heat map analysis showing expression of genes involved in building a peptidoglycan precursor Lipid II (D), Lipid II flippase and peptidoglycan polymerisation (E) and peptidoglycan degradation and recycling (F). G. RTqPCR data showing relative expression levels of 19 Ot genes involved in bacterial growth and division at 3-, 4-, 5- and 6-days post infection taken from 5 independent biological replicates. Gene expression is normalised to *infB, rpoB, rpoC* and *gyrB.* Statistical significance was determined using a two-way ANOVA followed by Tukey’s multiple comparisons. Statistical significance compared to 3 d.p.i is shown. ** p < 0.01, **** p < 0.0001.

Ot has been shown to encode an unusual peptidoglycan-like structure that is particularly low in abundance and built in the absence of the usually essential peptidoglycan polymerization protein class A PBP^35,63,64^. However, Ot encodes the main components of the divisome and elongasome, which function to assemble the peptidoglycan polymerization machinery (Fig. 5A). We analyzed the relative expression levels of elongasome and divisome genes across the strains in our study (Fig. 5B). We then determined the relative levels of genes directly involved in assembling the peptidoglycan components (Fig. 5C). We analyzed three distinct groups: (i) those involved in assembling the peptidoglycan precursor lipid II, (ii) those involved in flipping lipid II to the periplasmic side and polymerizing it into elongating peptidoglycan, and (iii) those involved in degradation and recycling (Fig. 5D-F). Together, the analysis showed that the elongasome/divisome genes, as well as those directly involved in peptidoglycan synthesis and turnover, exhibit some variation between strains. This may reflect differences in the activities of these pathways, or the fact that the pathways are differentially utilized at different stages of the intracellular infection cycle. Small differences in the growth of different strains would mean that the samples collected for RNA sequencing analysis would be at slightly different stages of the intracellular life cycle. To further probe the variation in gene expression over time, we collected RNA from one Ot strain, Karp, at 3-, 4-, 5- and 6-days post infection and measured the relative expression of selected elongasome/divisome and peptidoglycan biosynthesis genes at each time point. We also included *scaA* to compare these data with previous analyses^19^ and confirmed that expression of this transcript increased over time although this diference was not statistically significant (Fig. 5G). Two genes exhibited large and statistically significant increases in expression over time: the UDP-N-acetylenolpyruvoylglucosamine reductase *murB,* and the lytic transglycolsylase *slt.* This does not correspond with strain-to-strain transcript variability in the RNAseq dataset because *murB* exhibits low variability, whilst *slt* exhibits higher transcript variability but lower changes over time. Together these data show that certain peptidoglycan biosynthesis genes are transcriptionally regulated through the Ot intracellular infection cycle.

## Discussion

In this study, we compared the transcriptional landscapes of eight Ot strains during infection of human endothelial cells. By analyzing both host and bacterial transcriptomes, we identified pathways consistently activated across strains as well as those with strain-specific patterns of regulation.

A central finding is that all strains elicit a robust type I interferon response in endothelial cells, underscoring the highly conserved host recognition of Ot. However, the downstream gene expression profiles varied markedly between strains, independent of virulence or phylogeny. Notably, Gilliam showed reduced induction of multiple pattern recognition receptors, particularly NOD2, while TA686 uniquely upregulated MHC-II–associated genes, in contrast to their downregulation by other strains. TA686 and UT176 further altered expression of cell cycle genes, whereas Gilliam disproportionately impacted translation-related pathways. These results support a model in which each strain generates a distinct “immunological fingerprint,” with relatively subtle transcriptional differences that, when integrated across the immune system in vivo, may contribute to divergent disease outcomes.

On the bacterial side, we observed no global transcriptome patterns correlating with virulence, phylogeny, or host response, reinforcing earlier findings that virulence cannot be explained by single genetic determinants alone. Instead, the most striking variation lay in the expression and repertoire of effector proteins, particularly the large and heterogeneous Ankyrin repeat family. The specific complement of Anks expressed by each strain likely underpins differential modulation of host pathways, contributing to the unique strain-dependent signatures of host response. One example of this is in the modulation of host cell cycle. It was shown that Ot Ikeda downregulates expression of the cell cycle checkpoint regulator p53 through Ank13 activity^38^. This particular Ank is not conserved in all Ot strains and is absent from strain TA686 which we show here exhibits a different pattern of cell cycle. This likely reflects distinct modes of host cell modulation driven by the specific Ankyrin repertoires of individual strains.

We also investigated the regulation of bacterial growth and cell wall pathways, focusing on the divisome, elongasome, and peptidoglycan biosynthesis machinery. Most components of these pathways showed relatively stable expression across strains, suggesting constitutive activity during intracellular replication. However, temporal analysis revealed that *murB* and *slt* transcripts increased significantly between days 3 and 6 post-infection. These enzymes act at the initiating and terminal steps of peptidoglycan metabolism, respectively, suggesting that they may represent regulatory checkpoints in the synthesis and turnover of Ot’s reduced peptidoglycan-like structure. This is consistent with the proposed developmental cycle of Ot, in which cell wall remodeling accompanies transitions between intracellular and extracellular states^19^.

Taken together, our results highlight that while Ot strains uniformly engage innate immune signaling, the downstream host transcriptional profiles are uniquely tuned by each strain’s effector repertoire. In parallel, bacterial growth and cell wall genes are broadly constitutive, with only select enzymes showing transcriptional regulation, pointing to potential control nodes in the developmental cycle.

This work provides the most comprehensive transcriptional comparison of Ot strains to date. By integrating host and bacterial responses, we show that variation in disease outcome cannot be attributed to a single gene or pathway, but rather emerges from the combined effects of diverse bacterial effectors acting on host signaling networks. The identification of strain-specific host response fingerprints and putative regulatory points in bacterial growth pathways opens new avenues for mechanistic studies. Ultimately, understanding how different Ot strains fine-tune host–pathogen interactions will be critical for unraveling the determinants of virulence and for guiding strategies to predict, diagnose, and control scrub typhus.

## Methods

### Mammalian cell culture

Mouse fibroblast L929 cell line (CCL-1, ATCC, USA) and Human Primary Umbilical Vein Endothelial Cells (HUVEC; PCS-100-010, ATCC, USA) purchased from the American Type Culture Collection) were cultured and used to propagate Ot for preparation of bacterial frozen stock, and for isolation of total RNA from infected cells, respectively. Mycoplasma Detection Kit (13100-01, Southern Biotechnology, USA), a PCR-based mycoplasma testing was applied to confirm that both the L929 cells and HUVECs were mycoplasma-free. The growth medium with supplement and growth condition are as indicated: L929 – Gibco™ High Glucose Dulbecco’s Modified Eagle Medium (DMEM; 11-965-092, Thermo Fisher Scientific, USA) supplemented with 10% heat-inactivated Fetal Bovine Serum (HI FBS; 16140089, Thermo Fisher Scientific, USA), at 37°C with 5% CO_2_. HUVEC – Gibco™ Human Large Vessel Endothelial Cell Basal Medium, formerly Medium 200 (M200500, Thermo Fisher Scientific, USA) topped up with 50X Gibco™ Large Vessel Endothelial Supplement (LVES; A1460801, Thermo Fisher Scientific, USA), at 37°C with 5% CO_2_. The host cells were grown in either a Nunc™ T25 or T75 culture flask (156367 and 156499, respectively, Thermo Fisher Scientific, Denmark) to 100% confluency and subcultured up to 10 passages to yield 80% confluency for 24 hours before bacterial propagation following a standard cell culture procedure.

### Bacterial strains, propagation, DNA extraction, and quantification

Seven strains of *Orientia tsutsugamushi* consisting of Ikeda, Karp, TA686, UT76, TA763, Gilliam, and Kato were used in this study (Supp. Table 1) and grown as described previously^65^. The genome sequences and source of the strains is described here^25^. The bacteria were propagated routinely in either a T25 or T75 culture flask pre-seeded with a monolayer of L929 cells in DMEM supplemented with 10% HI FBS grown at 37°C with 5% CO_2_. We then harvested intracellular Ot and extracted the bacterial DNA using an alkaline lysis buffer^65^. A Bio-Rad CFX Connect Real-Time PCR Detection System Thermal Cycler (1855201 Bio-Rad, USA) and the primer/probe set were applied to amplify the bacterial single-copy gene *tsa47*, including tsa47F 5′-TCCAGAATTAAATGAGAATTTAGGAC-3′, tsa47R 5′-TTAGTAATTACATCTCCAGGAGCAA-3′ and tsa47probe 5′-[6FAM] TTCCACATTGTGCTGCAGATCCTTC[TAM]-3′ (Sigma-Aldrich, USA). To assess the quantity of Ot by determining genome copy numbers relative to a standard curve, we used the Bio-Rad CFX Manager software version 3.1.

### Total RNA extraction and quantification

The bacterial frozen stocks of each Ot strain were made from infected L929 cells and stored in 218mM sucrose phosphate glutamine buffer (SPG; 218mM sucrose, 3.76 mM KH_2_PO4, 7.1 mM KH_2_PO4, 4.9 mM monosodium L-glutamic acid) using the same procedure previously described^65^. We thawed the frozen stocks to inoculate a monolayer of HUVECs in three T25 flasks per strain to establish biological triplicate of pre-growth or the first passage. The intracellular fraction of pre-grown bacteria was then harvested and isolated from host cells on day 7 post-infection using the bead mill homogenization method^65^. Subsequently, the bacterial suspension of individual replicates from each strain was subject to *i.* alkaline lysis DNA extraction for *tsa47* qPCR to determine bacterial copy number as stated above, and *ii.* infection of 80% confluent HUVECs in a T25 flask for the second passage with the beginning amount and approximate MOI, as shown in Supp. Table 2. At six days post-infection, we harvested a total of 21 flasks of infected HUVECs from 7 strains in triplicates, starting by removing the media, washing them twice with sterile Gibco™ PBS (10010031, Thermo Fisher Scientific, USA), and disassociating the cells from the surface by Gibco™ 0.25% Trypsin-EDTA solution (25200056, Thermo Fisher Scientific, USA) with gentle manual agitation. Trypsin was then deactivated by mixing thoroughly with DMEM plus 10% HI FBS. 20-100 µl of this mixture was taken for DNA extraction and qPCR to determine the final number of Ot (Supp.Table 2) prior to RNA extraction. The remaining Ot-infected cell suspension was then centrifuged at 1000 xg for 5 minutes, and the supernatant was discarded to obtain the cell pellet, which was subsequently utilized as the starting material for RNA isolation using the spin column method by RNeasy Mini Kit (74106, Qiagen, Germany). We extracted RNA according to the manufacturer’s protocol with minor adjustments. We additionally eliminated the genomic DNA by performing on-column DNase treatment for 1 hour at room temperature using an RNase-Free DNase Set (79254, Qiagen, Germany) and proceeded to washing steps as usual. We eluted the RNA from the column with 30 µL of RNase-free water supplied in the kit. As demonstrated in Supp. Table 2, we exploited NanoDrop™ 2000 Spectrophotometers (ND-2000, Thermo Fisher Scientific, USA) to measure the quantity and quality of the 21 purified RNA samples, which were later shipped to the Core Unit SysMed, University of Würzburg in Würzburg, Germany for RNA sequencing.

### RNA processing and sequencing

Library preparation and sequencing was performed by the Core Unit SysMed Würzburg. RNA integrity was evaluated using a Bioanalyzer (Agilent). Ribosomal RNA was depleted using the RiboCOP (Lexogen) META and Human/Mouse/Rat kits according to the manufacturer’s instructions. Libraries were prepared using the Lexogen Corall Total RNA Library Prep kit. Sequencing was performed on a NextSeq 2000 platform (Illumina) at the Core Unit SysMed in single-end mode for 100 cycles.

### Bacterial annotations

Orientia genomes were downloaded from NCBI using the following accessions: Karp (GCA_900327275.1), Kato (GCA_900327265.1), Gilliam (GCA_900327245.1), Ikeda (GCA_000010205.1), UT176 (GCA_900327235.1), UT76 (GCA_900327255.1), TA763 (GCA_900327225.1) and TA686 (GCF_900377405.1). Additional bacterial gene annotations including for *5S*, *tmRNA* and *RnaseP* identified from Mika-Gospodorz et al^21^ were added to Karp and UT176. BLAST was used to identify these additional genes in other. The gene coordinates for these additional genes are available in (Supp. Table 4).

### Orientia gene characterisation

Annotations of Ankyrin, T4SS and TPR genes were retrieved from our previous annotation for all strains excluding TA763^24^. Ankyrin, TPR and T4SS genes were manually identified for TA763 from the genome annotation. Annotations for peptidoglycan genes were retrieved from Otten et al^64^.

### Orthologous bacterial genes

We used POFF^66^ to identify orthologous genes across the 8 strains. Genes were required to be present across all strains as a single copy. Gene sequences from the Karp genome annotation served as a reference for annotation with eggNOG-mapper^67^.

### Phylogeny

The bacterial phylogenetic tree was reconstructed from 706 orthologous genes identified above. Each orthologous gene was aligned across strains using MAFFT^68^ then AMAS was used to concatenate all alignments^69^. Phylogenetic inference was performed with RAxML-NG^70^ using the General Time Reversible (GTR) model with gamma-distributed rate heterogeneity (GTR+G). The maximum likelihood (ML) tree was inferred from the concatenated alignment and bootstrap values were calculated using 1,000 replicates.

### RNA-seq read processing and quantification

Raw sequencing reads were processed using the dualrnaseq pipeline https://nf-co.re/dualrnaseq/1.0.0, using BBDuk and Salmon selective alignment with additional parameters of “*--validateMappings --softclip --maxSoftclipFraction 0 0.8*“. Reads were then simultaneously mapped against the human (GRCh38) genome reference and the appropriate Orientia reference. TPM values were separately calculated for host reads and bacterial reads in R.

To classify the expression patterns of orthologous bacterial genes, we calculated mean TPM values using the three replicates for each strain^21^. Highly expressed genes had a mean TPM > 50, and expressed genes had a mean TPM > 10 (across all strains).

### Differential gene expression

Quantified reads were separated into host or bacterial reads and separately batch corrected with RUV-seq^71^, using RUVs with K=1. Differentially expression analysis was performed using edgeR^72^. Host genes were compared to mock-infected samples, using robust quasi-likelihood estimation and a false discovery rate (FDR) cutoff < 0.05.

### Highly variable bacterial genes

To quantify variability in gene expression across strains, we developed a method based on LOESS regression, loosely based on the FindVariableFeatures function in Seurat^73^. Briefly, a local regression curve was fitted to the variance across log_2_ mean normalized counts, and then the residual variance for each gene was calculated as the difference between the actual variance and that predicted by the regression curve. As expected, most genes had a residual variance around zero (Figure 3A). To identify genes with excess variability, we set a conservative threshold of greater than three times the median absolute deviation of the residual variance. For genes with low variability, we selected those with a LOESS residual of less than -0.25.

### Functional pathways and enrichment

Clusterprofiler was used for host KEGG enrichment (FDR < 0.05) and subsequent dotplots in Figure 2A^74^. Complexheatmap^75^ was used for all heatmaps. Host pathways for the functional analysis presented in Figure 2H were retrieved from InnateDB^76^ and loadings were compared between genes in each gene set with those outside it using a Wilcoxon rank-sum test for principal components 1 and 2 (Figure 1D).

### RTqPCR analysis

Ot_Karp bacteria were harvested from L929 cells at 3-, 4-, 5- and 6-days post-infection (dpi) and were stored in RNAProtect Bacteria Reagent (Qiagen 76506) at -80°C. Total RNA was extracted using the Qiagen RNeasy Plus Kit (Qiagen 74136), following the manufacturer’s instructions. Approximately 6 µg of extracted RNA was treated with TURBO DNase (ThermoFisher AM2238) at 37°C for 4 h. The amount of RNA used for reverse transcription in each sample was then calculated in relative to the Ot copy number in the 3-dpi sample. The RNA was reverse transcribed into cDNA using UltraScript^®^ 2.0 cDNA Synthesis Kit (PB30.32) and random hexamer. Controls lacking reverse transcriptase demonstrated that there was no contamination of genomic DNA in the RNA samples used for cDNA synthesis. Synthesised cDNA was diluted 1:10 and 1 µl was used as the template for qPCR using qPCRBIO SyGreen Mix Lo-ROX (PB20.11) with primers for each target gene (*ftsZ*, *mreB*, *ftsQ*, *murA*, *murB*, *murC*, *murD*, *murE*, *murF*, *murG*, *murJ*, *mraY*, *PBP2*, *rodA*, *ftsI*, *ftsW*, *PBP4*, *amiD*, *slt*, *scaA*, *infB*, *rpoB*, *rpoC*, *gyrB* and *16s*). Gene expression levels were calculated using the 2^(-ΔΔCt) method and were normalised to an average of four genes: *infB, rpoB, rpoC* and *gyrB*.

### Flow cytometry

Human umbilical vein endothelial cells (HUVECs) were cultured overnight in M200-500 medium (Gibco) supplemented with LVES (Gibco) and Penicillin–Streptomycin (Sigma-Aldrich) at 37 °C with 5% CO₂. Cells were infected with Ot at a multiplicity of infection (MOI) of 100. Following a 24-h incubation, cultures were washed twice with unsupplemented medium and further incubated for 5 days in growth medium.

Cells were detached using 0.25% trypsin (Gibco) and washed once with PBS. Unless otherwise stated, all centrifugation steps were performed at 500 × g for 5 min. Live/dead discrimination was performed using Zombie Red (BioLegend; 1:500 in PBS) for 30 min on ice, followed by washing and surface staining with mouse anti–HLA-DRB1 (Invitrogen; 1 µg/mL in PBS + 1% BSA) for 30 min on ice. Cells were fixed and permeabilised in ice-cold 70% ethanol overnight at 4 °C, washed with PBS, and intracellular staining was carried out in PBS + 1% BSA + 0.1% saponin using in-house human anti-TSA56 antibodies at 4 °C overnight. After three PBS washes, secondary antibodies (anti-human AF488 and anti-mouse AF647; Invitrogen; 1:200) were added and incubated for 1 h on ice in the dark. Cells were washed three times with PBS and stained with Hoechst 33342 (1 µg/mL) for 30 min. Samples were washed once and acquired on a BD LSRFortessa. Cell-cycle distributions (%M, %G1, %S, %G2/M) were quantified on live, singlet, Hoechst-positive populations using FlowJo v10.

## Data availability

Sequence reads are available on the sequence read archive (SRA) under accession number PRJNA1047511

## Acknowledgements

This work was funded by a Wellcome Trust Senior Research Fellowship (224277/Z/21/Z) to JS and a NSERC Discovery Grant (RGPIN-2024-04305) to LB. We additionally acknowledge compute resources provided by bayresq.net, funding by the Bavarian State Ministry for Science and Art, and the Core Unit SysMed at the University of Würzburg for sequencing services.

## Competing interests

The authors declare no competing interests.

## Notes

### Competing Interest Statement

The authors have declared no competing interest.

