## Supplementary Legends for "Comparative Dual RNA-Seq Analysis of Eight *Orientia tsutsugamushi* Strains in Endothelial Cells Reveals Regulatory Patterns in Bacterial and Host Pathways During Infection"

### Supplementary Tables

#### Supplementary Table 1

Strains of *Orientia tsutsugamushi* used in this study.

#### Supplementary Table 2

Information of the samples, pre- and post-RNA isolation

#### Supplementary Table 3

Number of sequenced reads from each biological sample.

#### Supplementary Table 4

RNAseq data on bacterial genes.

**Bacterial orthologs:** Ortholog table containing locus tags for each ortholog detected using POFF.

**Bacterial counts for orthologs:** Raw RNA-seq read counts for orthologous genes in each strain and replicate.

**Loess residuals:** Residual after fitting a curve to the mean-variance relationship in the bacterial data for each gene.

**Bacteria_STRAIN_T4SS**: Read counts for all type 4 secretion system-related genes in each strain.

**Bacteria_STRAIN_ANKs:** Read counts for all Ankyrin genes in each strain.

#### Supplementary Table 5

RNAseq data on host genes.

**Merged_DEGs:** Merged differential expression results for host cells infection with each strain. ENSG: Ensembl gene identifier; gene_name: common gene name; STRAIN_logFC: logFC for cells infected with STRAIN compared to mock infected; STRAIN_logCPM: average log counts per million for the comparison; STRAIN_F: F-statistic for the comparison; STRAIN_Pvalue: P-value for the comparison; STRAIN_FDR: FDR-corrected P-value for the comparison.

**InnateDB_PCX_enrichment**: Comparison of loadings values for genes in a selected gene set, compared to genes outside the geneset. SetSize: size of gene set; Median_in_set: median loading inside the gene set; Median_out_set: median loading outside the gene set; p_value: P-value for a Wilocoxon rank-sum test on the signed loadings; FDR: FDR-corrected p-values.

#### Supplementary Table 6

List of primers used in RTqPCR analysis of peptidoglycan-related genes.

### Supplementary Figures

#### Supplementary Figure 1

Heatmaps showing the log_2_ fold-changes (logFC) between infected and mock-infected cell for genes encoding pattern recognition receptors.

#### Supplementary Figure 2

Heatmaps showing the log_2_ fold-changes (logFC) between infected and mock-infected cell for interferon stimulated genes.

#### Supplementary Figure 3

Heatmaps showing the log_2_ fold-changes (logFC) between infected and mock-infected cell for IRF3/7 responsive genes.

#### Supplementary Figure 4

Heatmaps showing the log_2_ fold-changes (logFC) between infected and mock-infected cell for NFkB responsive genes.

#### Supplementary Figure 5

#### A Wilcoxon rank-sum test was used to test the signed loadings within each gene set to those outside it for a difference in median value. Axes show the –log_10_ P-value from the test, while bubble sizes show the gene set size.

#### Supplementary Figure 6

Heatmaps showing the log_2_ fold-changes (logFC) between infected and mock-infected cell for genes associated with Cyclin A/B1 associated events during G2/M transition.

#### Supplementary Figure 7

Boxplot showing the expression levels of *vir-*T4SS and non-*vir-*T4SS genes in Ot. The non-*vir* genes primarily belong to the F-type T4SS of Ot but cannot all be definitively classified as such due to duplicated genes shared between *vir-* and non-*vir-*T4SSs. All duplicated genes are excluded from this plot.
