## Supplementary figures and images for "Comparative Dual RNA-Seq Analysis of Eight *Orientia tsutsugamushi* Strains in Endothelial Cells Reveals Regulatory Patterns in Bacterial and Host Pathways During Infection"

### Supp. Fig. 1

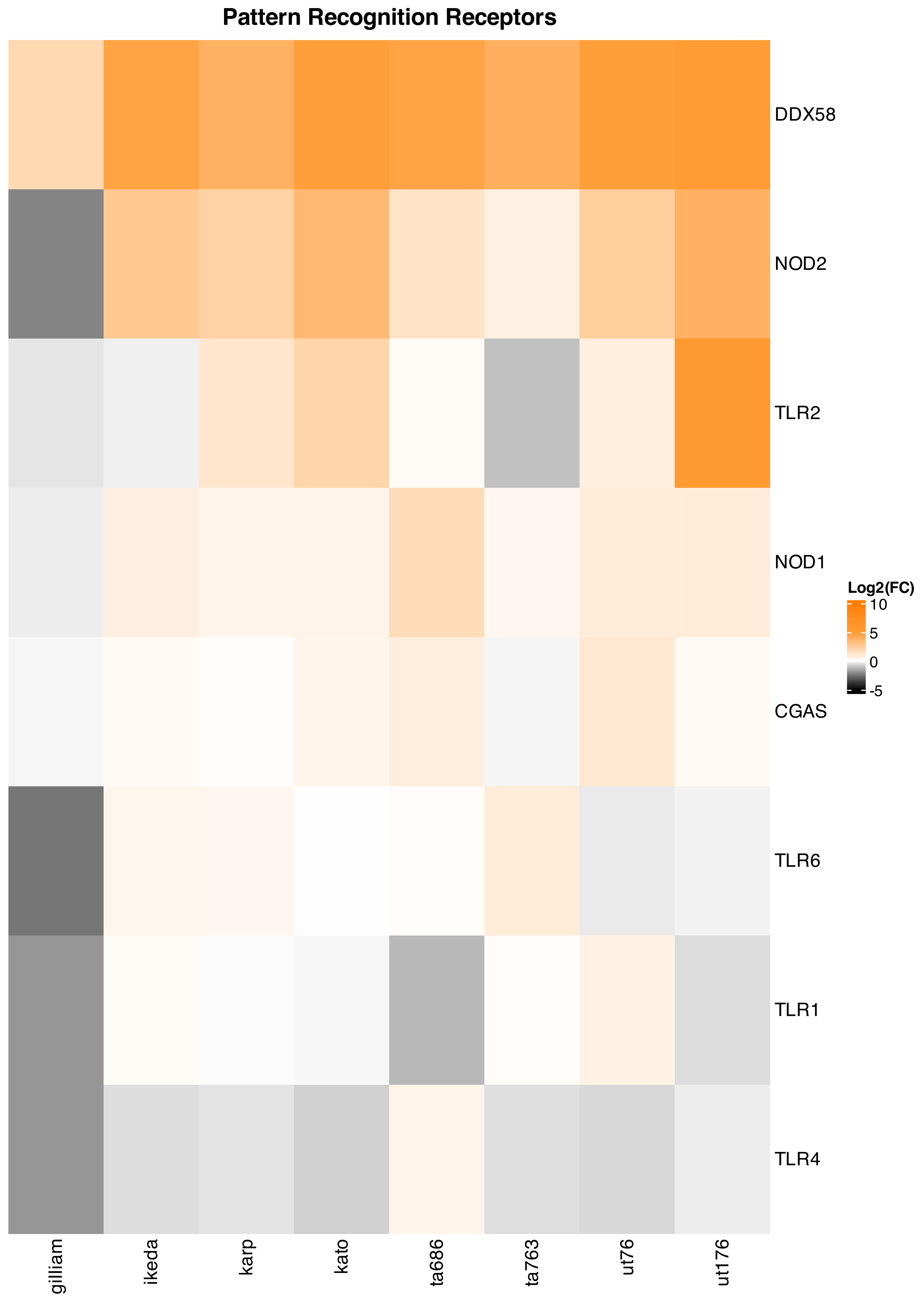

### Supp. Fig. 2

#### Interferon-stimulated Genes

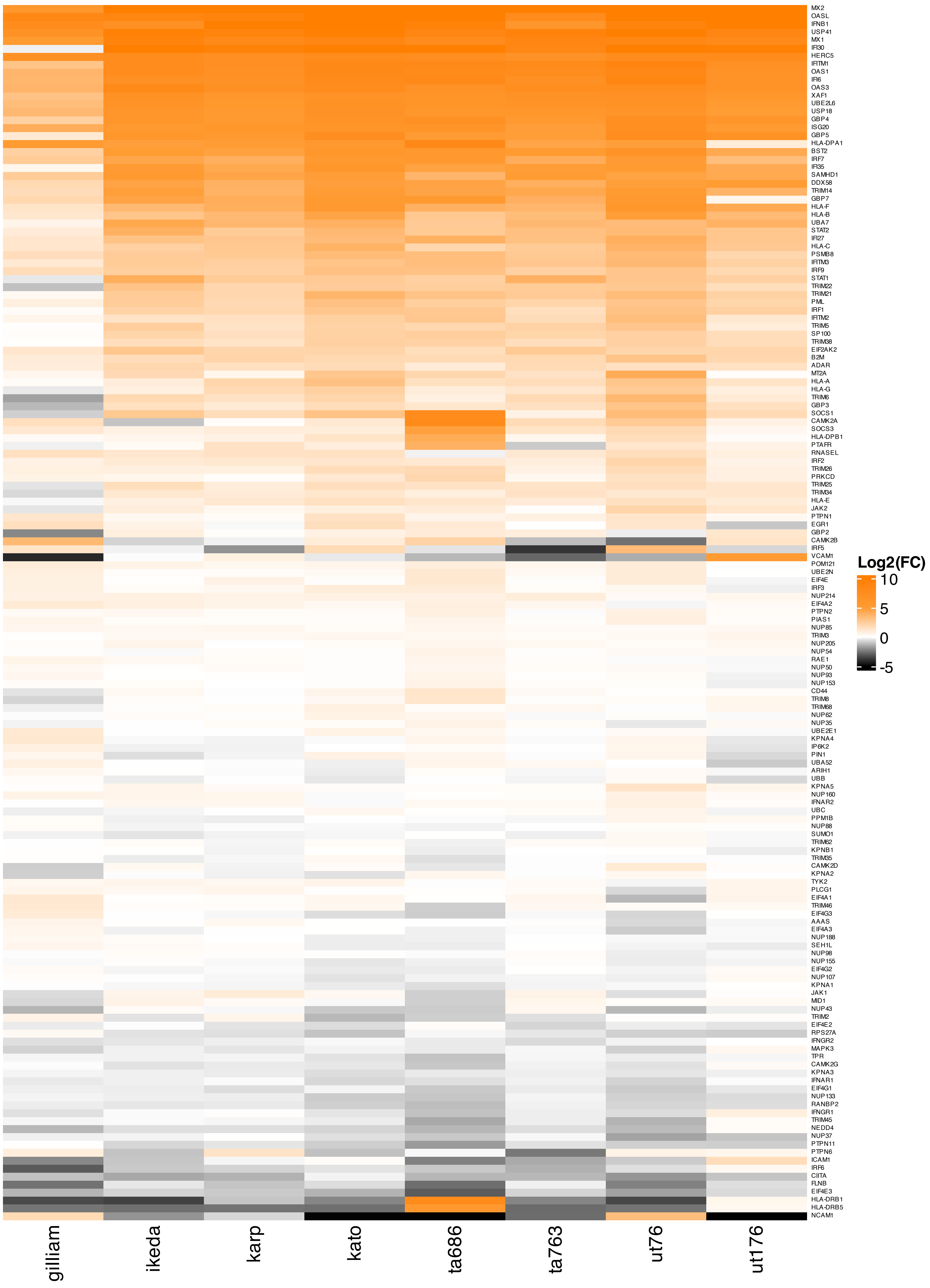

### Supp. Fig. 3

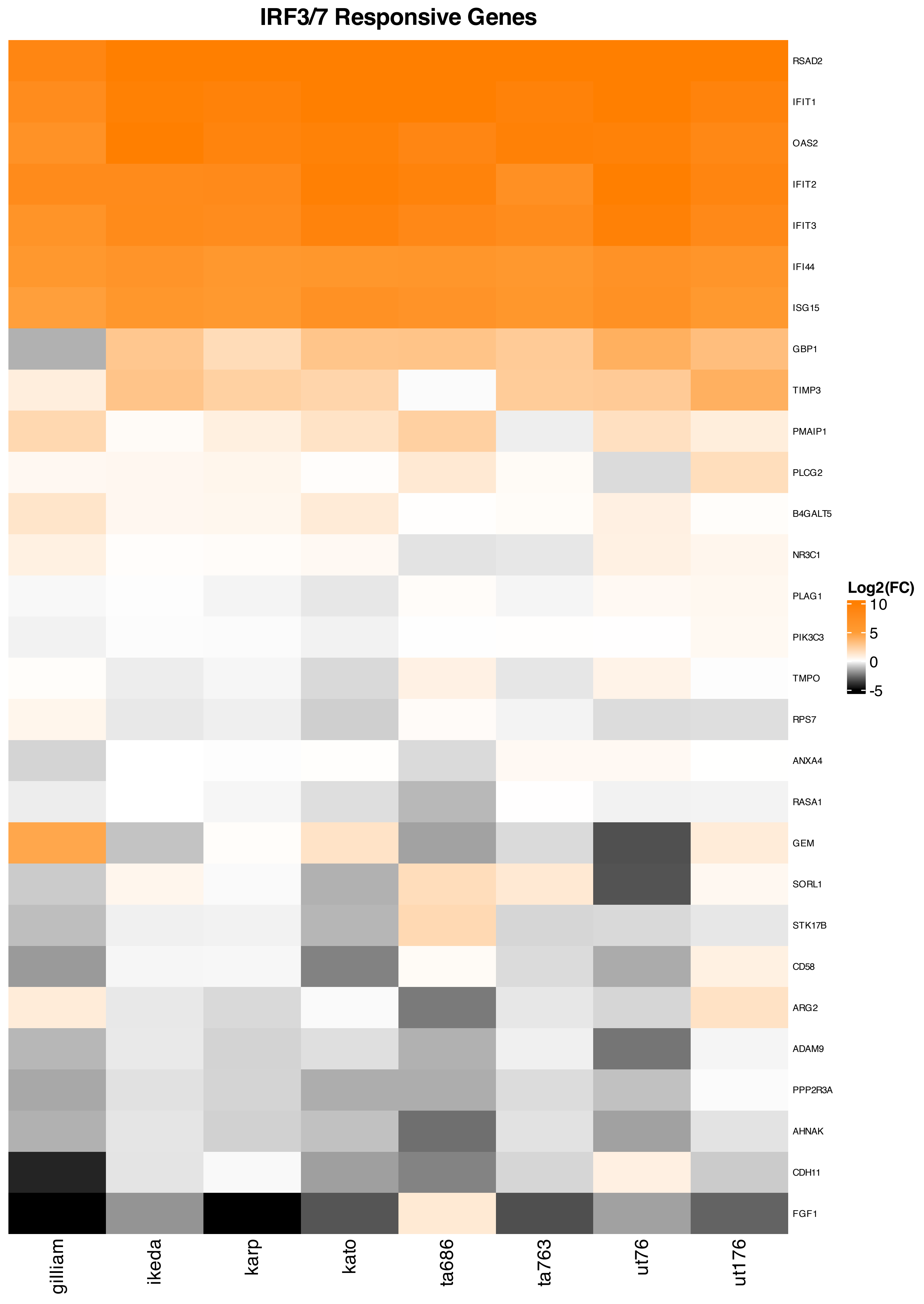

### Supp. Fig. 4

#### NFκB-dependent Genes

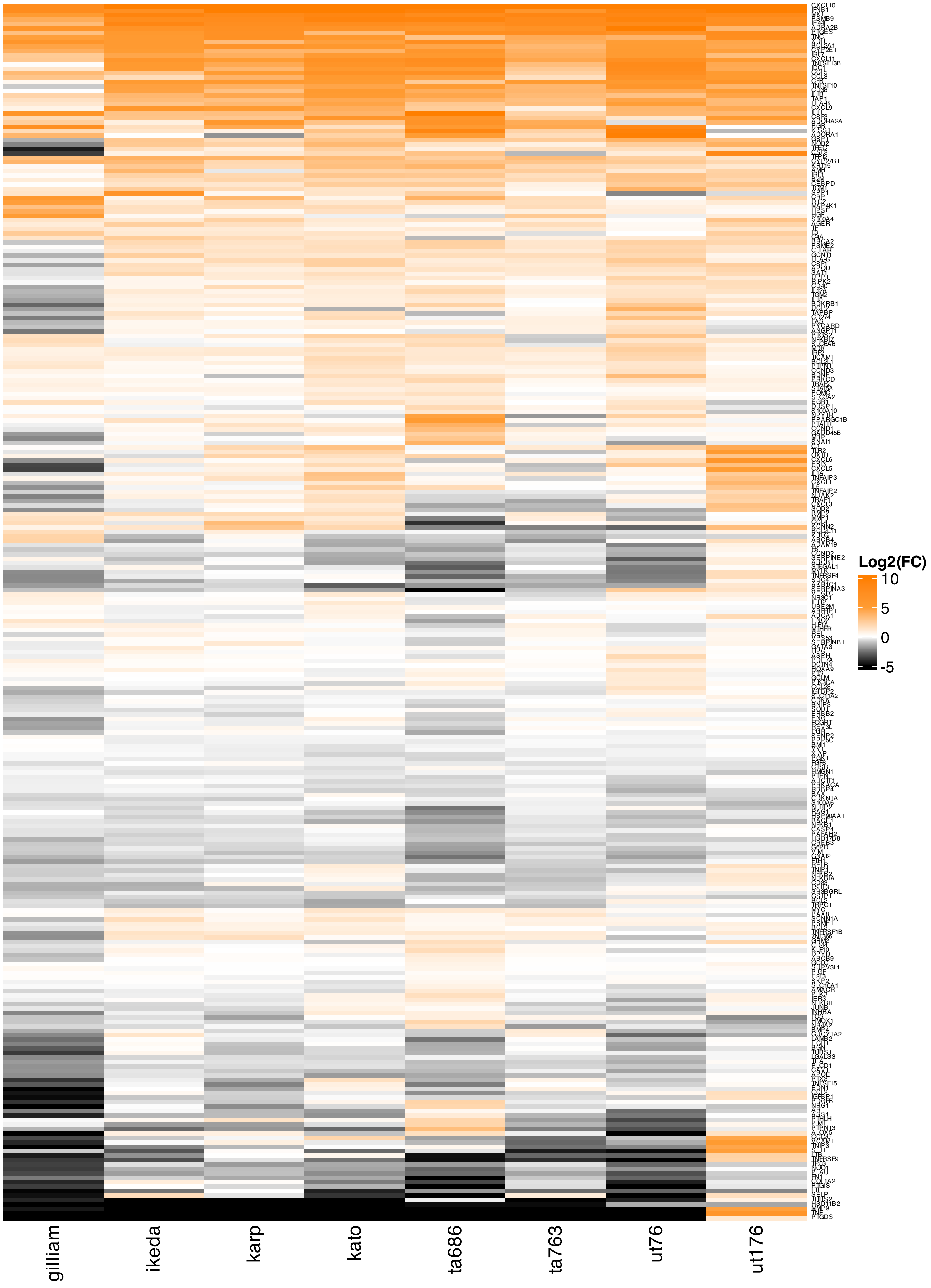

### Supp. Fig. 5

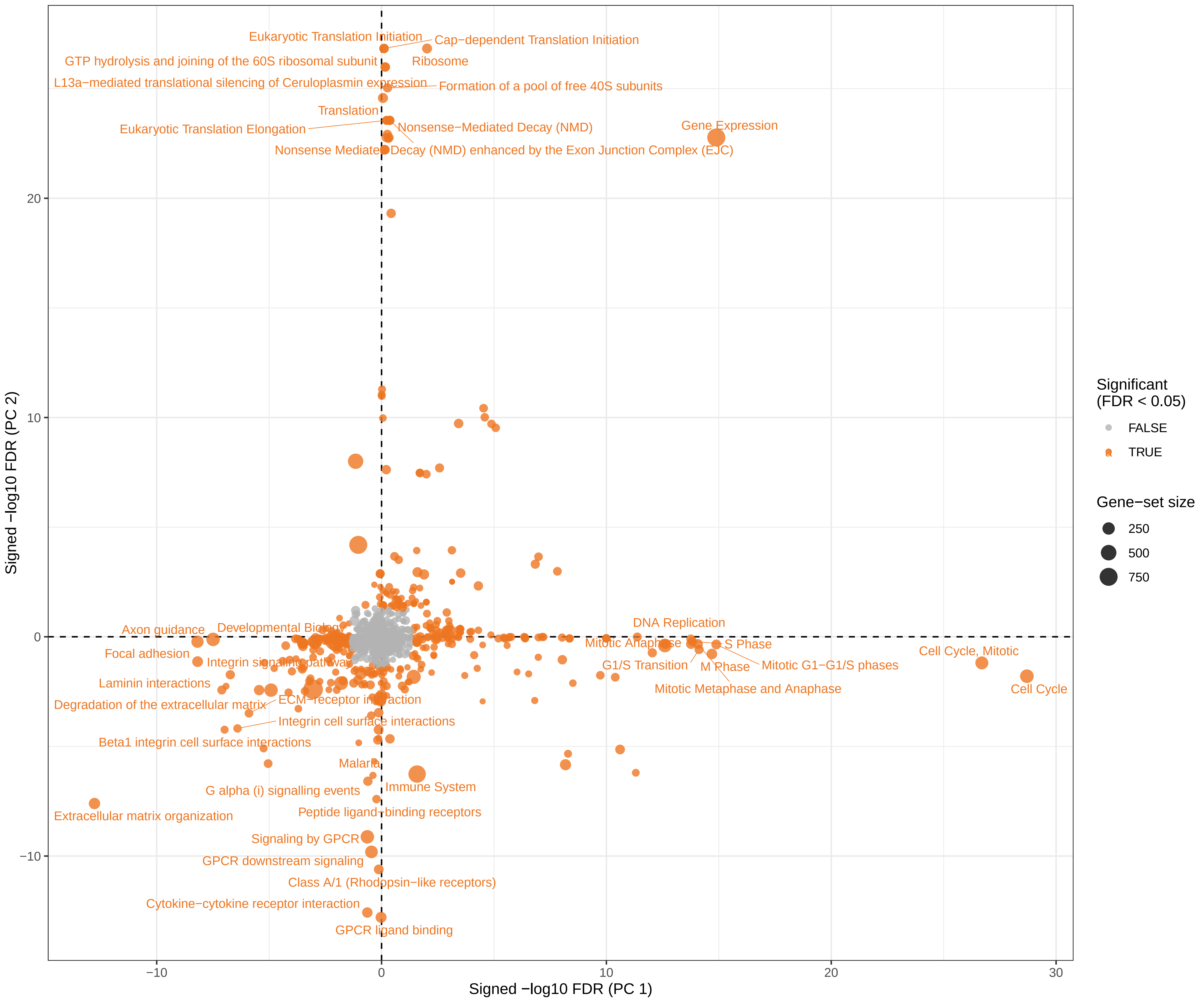

### Supp. Fig. 6

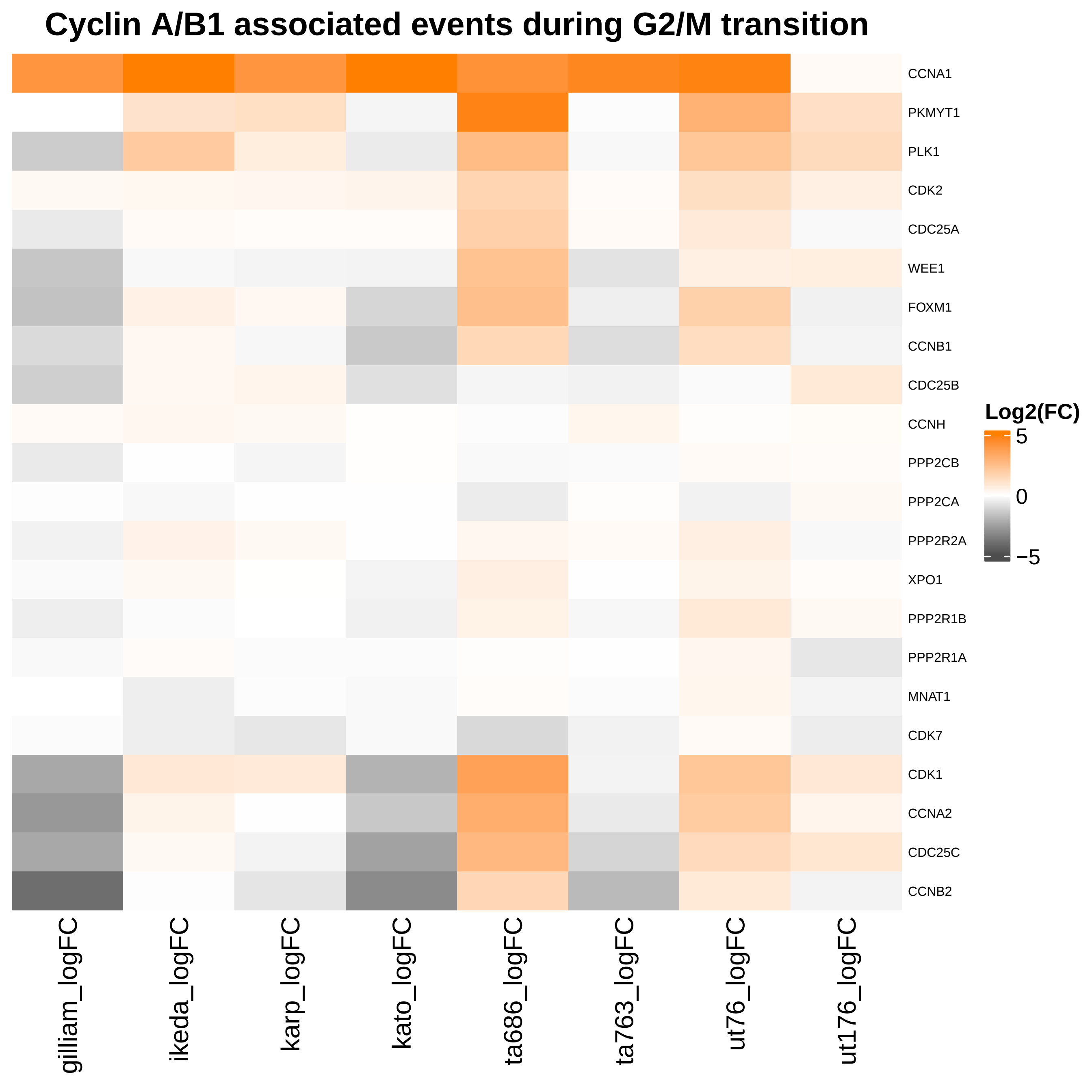

### Supp. Fig. 7

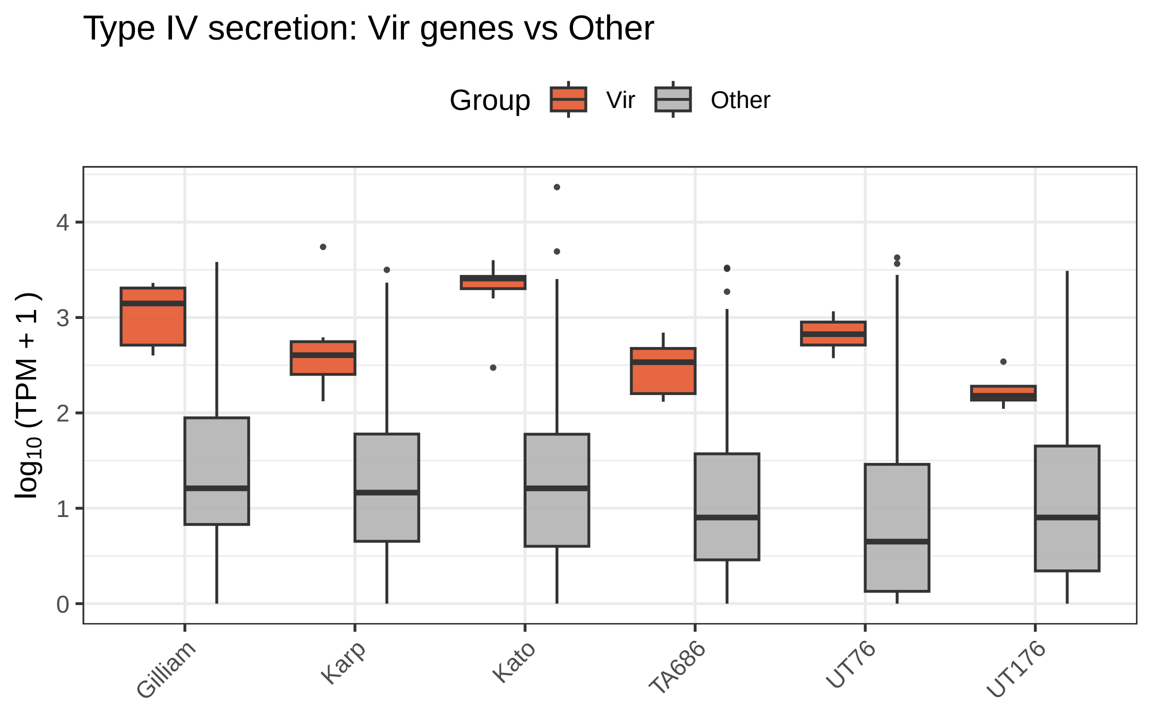
